# Experimental Assessment of Iris Water Content and Sex Differences in Mature Rabbits

**DOI:** 10.64898/2026.09.04.749502

**Authors:** Nima Rafezi, Sunaya A. Rahman, Babak N. Safa

**Author notes:** **Corresponding author:** 4202 E. Fowler Avenue, Tampa, FL 33620, USA.

## Abstract

Iris biomechanics plays a key role in primary angle-closure glaucoma (PACG). Soft tissue biomechanics is heavily influenced by tissue water content; therefore, accurately assessing water content is needed to describe iris biomechanics in a physiologically relevant manner. Yet, accurate experimental measurements of iris water content are rare in the literature. Therefore, our objective was to experimentally assess iris water content and potential sex differences in New Zealand White rabbit iris—a suitable model for studying iris biomechanics. We also examined the effect of exposure to phosphate-buffered saline (PBS) on iris water content, given its common use to maintain hydration during soft-tissue biomechanical testing. Our results showed slightly higher water content in males (89.2% ± 2.3%) than in females (87.7% ± 2.2%) (p = 0.022), while we found no significant differences in gross iris morphology between the sexes, indicating subtle compositional differences. Furthermore, PBS incubation had a minimal effect on water content only in female irides. Overall, our results indicated high iris water content in both sexes compared with anatomically adjacent ocular tissues and other soft connective tissues. These results improve our understanding of iris composition and physiology and can serve as a baseline for constructing biomechanical models to assess its role in PACG.

## Introduction

Primary angle-closure glaucoma (PACG) is a major subtype of glaucoma, the leading cause of irreversible blindness (Jonas *et al*., 2017; World Health Organization, 2019) where occlusion of the outflow pathway with the iris leads to intraocular pressure (IOP) elevation (Amerasinghe and Aung, 2008). PACG has a higher prevalence among women and people of Asian descent (Cheng *et al*., 2014; Sun *et al*., 2017). Although there are known physiological risk factors that connect certain anatomical parameters, such as anterior chamber depth and angle, to PACG (Nongpiur *et al*., 2011; Wang *et al*., 2023) there are no robust patient-specific risk stratification criteria for PACG (He *et al*., 2019; Yuan *et al*., 2023). Therefore, a deeper understanding of the iris and PACG is desired.

Iris biomechanics is closely associated with PACG (Safa *et al*., 2022; Mapstone, 1968; Tan *et al*., 2024). Notably, modern *in vivo* anterior segment optical coherence tomography (AS-OCT) combined with inverse FEM has improved the chances of clinical deployment of biomechanics for PACG risk assessment. As such, recent studies have suggested that human irides with PACG history tend to have a higher relative stiffness and lower hydraulic permeability compared to healthy eyes (Pant *et al*., 2018; Panda *et al*., 2021; Sebastian *et al*., 2025). Although these results have been partially corroborated by experimental tests such as AFM studies (Narayanaswamy *et al*., 2019), the *in vivo* studies often lack direct experimental validation and input parameters, such as soft tissue water content, needed for physiologically relevant modeling.

Water content is a key factor that influences the biomechanical response of soft tissues (Shahmirzadi and Hsieh, 2010; Meyer, McAvoy and Jiang, 2013). Water content may also be implicated in glaucomatous changes in iris composition, such as altered glycosaminoglycan (GAG) levels that could potentially affect iris hydration, as previously reported (Knepper *et al*., 1996). Furthermore, in soft tissues, phosphate-buffered saline (PBS) exposure can increase water content and reduce tensile stiffness, depending on exposure duration and tissue type (Screen *et al*., 2006; Han *et al*., 2012; Safa *et al*., 2017). For iris, only a few studies have reported water content as a secondary measurement (Coben *et al*., 1970; Ellis, Littlejohn and Deitrich, 1972; Dutton *et al*., 1981; Stone and Wilson, 1982); however, they do not address potential confounding factors (e.g., sex differences) or lack important technical controls (e.g., accounting for potential postmortem changes and the effect of buffer solution during tissue handling).

In this study, our objective was to provide a detailed experimental assessment of iris water content and the associated sex differences in rabbit iris. We chose rabbits because their anterior chamber anatomy closely resembles the human eye, making them suitable for studying iris biomechanics (Peiffer, Pohm-Thorsen and Corcoran, 1994; Yamaji *et al*., 2003). Additionally, since postmortem pupillary dilation can impact iris volume, it is plausible that such changes can influence iris water content (Quigley *et al*., 2009); therefore, we also measured the relative state of postmortem pupil dilation as a control in our experiments. Furthermore, motivated by the higher prevalence of PACG in women (Vijaya *et al*., 2006; Cheng *et al*., 2013; Zhang *et al*., 2021), we treated sex as a biological variable to detect potential sex differences in iris water content. Finally, since the commonly used PBS buffer for biomechanical testing can affect the water content and the biomechanical properties of soft connective tissues, we also tested the effect of PBS exposure on iris water content. Overall, these measurements can improve understanding of iris physiology and support biomechanical modeling of the iris tissue to elucidate the association between biomechanics and PACG.

## Methods

### Animal Tissue

In this study, we obtained fresh eyes from fully mature New Zealand White rabbits (c. 6 months old) from a commercial vendor (Pel-Freez Biologicals, Rogers, AR, US). We followed ARVO’s guidelines on the use of animals in vision research (Association for Research in Vision and Ophthalmology, 2024). We received globes shipped in PBS on ice, and we conducted all tests within 1 day postmortem. We used 51 globes (25 males and 26 females). We performed all analyses on approximately equal numbers of male and female eyes, treating sex as a biological variable.

### Iridial Gross Morphology

Pupillary dilation and constriction can alter the volume of the iris (Mak, Xu and Leung, 2013) enabled by fluid exchange between the highly permeable iris stroma and the surrounding aqueous humor (Mark, 2003; Tan *et al*., 2019). A differential postmortem dilation ratio may confound iris water content measurements. To control for this effect, we conducted a gross morphology analysis. We captured frontal images of a subset of the globes (16 males and 17 females) and used Fiji (Schindelin et al., 2012) to annotate the limbal and pupillary margins with closed splines and measure the enclosed area (A) of each curve (Figure 1). Due to the semi-circular (slightly oval) shape of the rabbit pupil and limbus, we calculated the equivalent pupillary and limbal diameters as *d = √4A/π*, and the postmortem dilation ratio, which we defined as the ratio between equivalent pupil and limbus diameters. To address annotation subjectivity, three annotators independently marked the margins, and we used the average of the three for each eye in the final analysis (see Supplementary Document Figure S1 for further information).

**Figure 1:**
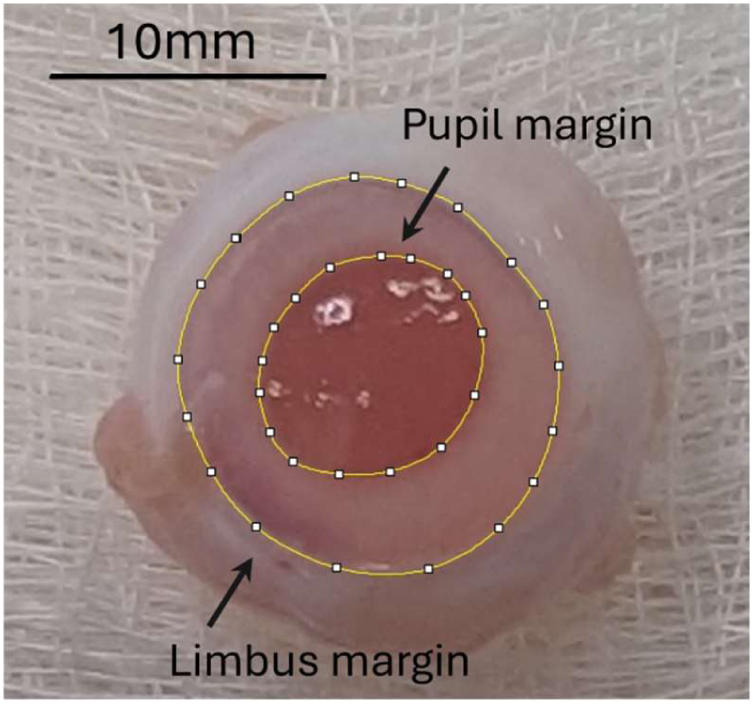
Frontal picture of the rabbit eye globe, demonstrating the limbus and pupil margins and their annotations.

### Baseline Water Content Measurement

Motivated by the connection between tissue hydration level and biomechanical response, and the need for tissue water content (i.e., ratio of water mass to total tissue mass) in computational analyses of iris biomechanics (Carniel *et al*., 2023; Shahmirzadi and Hsieh, 2010), we measured the water content of rabbit irides in all the eyes analyzed (*n* = 51; 25 male, 26 female). For these experiments, we used a portion of the iris (one-third to half), while the rest of the iris was either used for additional tests in this study (i.e., for the effect of the PBS experiment, explained below) or in an unrelated study aiming to minimize the need for additional animal sacrifice. Before mass measurement, we removed excess aqueous humor on the tissue surface by gently blotting the tissue with non-abrasive cellulose tissues (Kimwipes; Kimtech, Roswell, GA, US), then placed each sample in a pre-weighed microtube with the cap closed to minimize ambient dehydration.

We measured the total mass of the sample before drying (*m*_wet_), and then measured the solid weight (*m*_dry_), after drying the sample in an oven at 63°C to avoid collagen denaturation (Bozec and Odlyha, 2011). We used three to four days of drying on four similar batches of eyes (5-7 samples per sex), where each batch was measured over at least three days. We measured each sample three times in pre-weighed microtubes using an analytical balance (±0.1 mg) and averaged the measurements across days. The measured mass was several hundred times greater than the scale uncertainty, ensuring sufficient measurement sensitivity. Finally, we calculated the water content as φ = (*m*_wet_ − *m*_dry_)/*m*_wet_ (Safa *et al*., 2017).

### Effect of PBS Exposure on Water Content

Exposure to buffer solutions can alter water content and biomechanical properties. Because PBS is widely used in mechanical testing, we compared the water content of PBS-exposed samples with their paired untreated controls (6 males and 6 females). To mimic potential exposure time in a typical biomechanical test, we incubated the irides in PBS for 20 minutes. One half of the iris served as a control, and the other half was exposed to PBS at room temperature. We used the same protocol to measure water content for the baseline measurements explained above.

### Statistical Analysis

We used unpaired Welch’s t-test to compare male and female gross morphology results. To test whether our drying method was sufficient to achieve equilibration, we conducted a one-way ANOVA and post hoc Tukey’s multiple comparison tests on tissue mass normalized to the initial mass of the tissue (*m*_wet_) across the days of drying. Furthermore, we used unpaired Welch’s t-tests to compare baseline water content between male and female irides. Finally, to assess the effect of PBS exposure on water content, we used a two-factor repeated-measures ANOVA with sex (male vs. female) as the independent factor and treatment (control vs. PBS) as the paired factor, followed by a Bonferroni multiple-comparisons test. To support interpretation of significant t-tests, we also reported effect size by calculating Cohen’s d as the difference between means divided by the pooled standard deviation (Lakens, 2013). We set the significance level at 5% for all tests. We conducted these statistical analyses using GraphPad Prism (version 11.0.1 for Windows; GraphPad Software, Boston, MA, US, www.graphpad.com).

## Results

The assessment of the gross morphology of the irides before dissection indicated no difference in the diameter of the limbus (p = 0.385) and pupil (p = 0.447) between male and female irides (Figure 2). The average rabbit iris limbus and pupil diameters were 15.1 ± 0.9 mm (mean ± standard deviation) and 8.1 ± 1.1 mm, respectively (Figure 2A and B). Additionally, postmortem dilation did not differ between the sexes (p = 0.627), and the average ratio was 53.9% ± 6.1% (Figure 2C).

**Figure 2:**
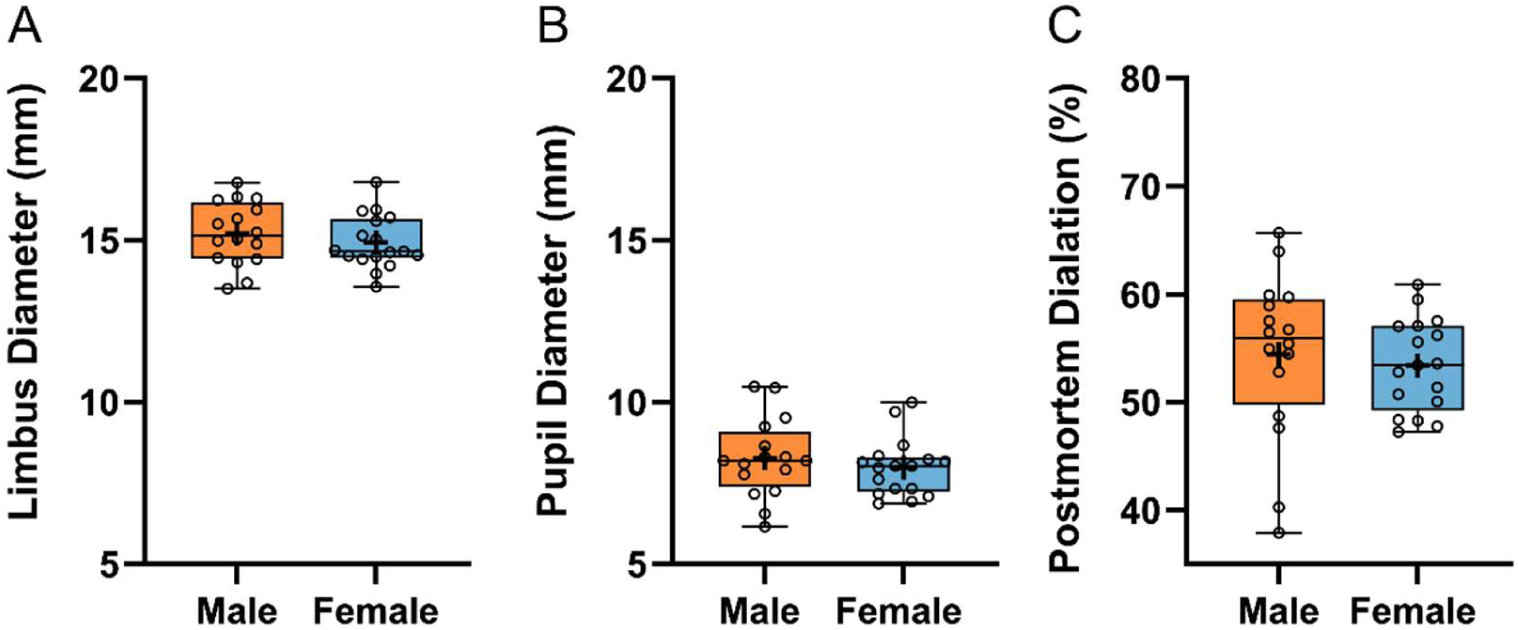
The results of gross morphological analysis, showing (A) the equivalent limbus diameter, (B) equivalent pupil diameter, and (C) the postmortem dilation ratio, calculated as the ratio of the equivalent pupil diameter to equivalent limbus diameter. Overall, we found no sex differences between samples on either measure (p>0.05). The individual data points are shown and overlaid with box plots including horizontal solid lines demonstrating the median and interquartile range (IQR). The mean value is shown with a ‘+’ symbol, and the whiskers indicate the minimum and maximum data values.

The samples achieved mass equilibration across the drying days tested (Figure 3). This trend was consistent for male (p = 0.060) and female (p = 0.815) samples. Similarly, post hoc multiple comparisons showed no significant differences across drying days for male (p > 0.119) and female (p > 0.846) samples. We noted some small variation in water content in male samples on day 4, which we attribute to experimental variability or reabsorption of ambient moisture rather than a physiological effect.

**Figure 3:**
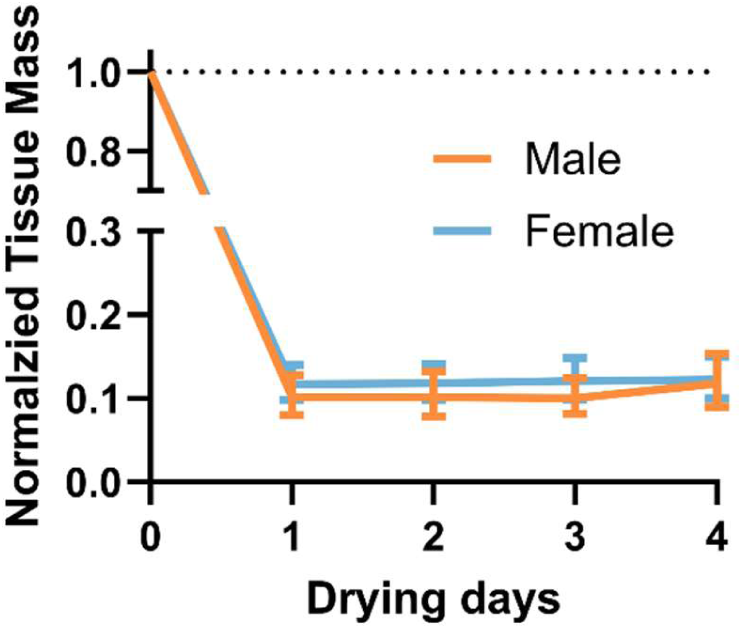
Normalized mass of iris samples over the course time, showing a drastic reduction in the weight of the samples after one day of oven drying, which was stable for up to four days. Data are shown as the mean, with standard deviation as error bars.

Interestingly, the water content of the male irides (89.2% ± 2.3%) was slightly higher than that of the females (87.7% ± 2.2%; d = 0.662; p = 0.022; Figure 4). Although this 1.5% mean difference in water content was statistically significant, it was smaller than the overall variation observed across all batches tested (pooled standard deviation = 2.3%).

**Figure 4:**
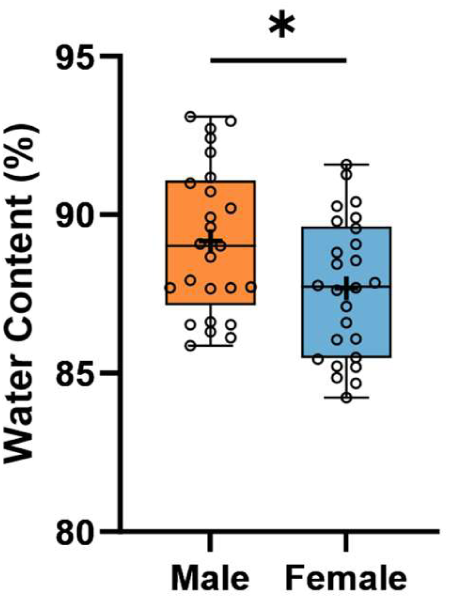
Iris water content calculated separately for the male and female samples, which indicated a modest (difference in the means = 1.5%), yet statistically significant (Cohen’s d = 0.662 and p = 0.022) higher water content in male samples. Data are shown as individual points with box and whiskers as before. *p < 0.05

Finally, exposure to PBS buffer for 20 minutes had a minimal effect on iris water content, despite some mixed statistical results (Figure 5). Based on a 2-way ANOVA test, while sex had no effect on this batch’s water content (p = 0.828), PBS treatment did (p = 0.014), and there was no interaction between these factors (p = 0.338). Subsequent multiple comparisons indicated no differences between sex groups (p > 0.943), and PBS exposure did not alter water content of the male sample (mean difference = 0.8%; d = 0.470; p = 0.394), whereas female samples showed a 1.6% mean difference (d = 2.275; p = 0.037). This difference was smaller than the overall variation across all batches tested (Figure 4).

**Figure 5:**
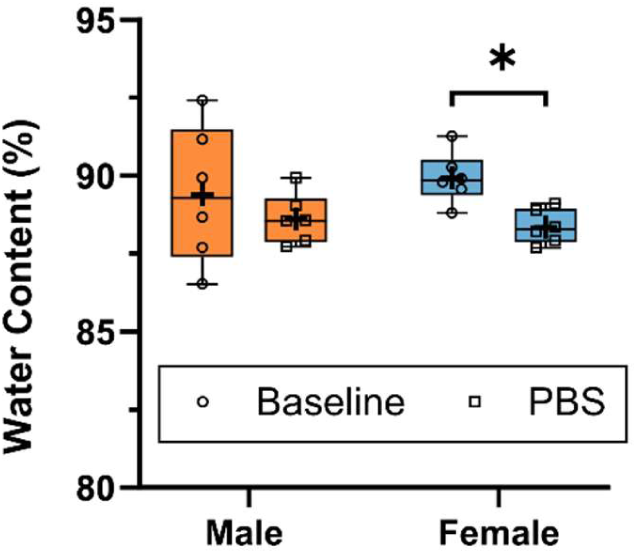
An analysis of the effect of PBS on the water content of both male and female samples indicated no effect on the male samples (Cohen’s d = 0.470 and p = 0.394), and a trivial difference in the female samples (Cohen’s d = 2.275 and p = 0.037). Data are shown as individual points with box and whiskers as before. *p < 0.05

## Discussion

Iris biomechanics contributes to the pathophysiology of PACG (Mapstone, 1968; Zheng *et al*., 2012; Panda *et al*., 2021; Tan *et al*., 2024). Motivated by the connection between soft-tissue (such as the iris) biomechanical properties and water content, we provided a detailed analysis of iris water content. More specifically, water content controls the poroelastic biomechanical behavior (fluid-dependent viscoelasticity) of soft tissues and shapes their response to dynamic mechanical loading. Because the iris controls the pupil, it experiences highly dynamic biomechanical loading; therefore, understanding its biomechanics requires understanding its water content. Some emerging studies have incorporated iris poroelasticity into their PACG analyses using FEM (Pant *et al*., 2018; Panda *et al*., 2021; Sebastian *et al*., 2025); however, experimental assessment of iris poroelasticity and water content remains incomplete. Our experimental results can serve as benchmark values to improve understanding of iris physiology and to develop computational models with improved physiological fidelity to better understand PACG pathophysiology.

Our analysis indicated that the iris has a high water content (c. 88%; Figure 4), which is generally in agreement with previous studies that reported c. 85% water content (Coben *et al*., 1970; Ellis, Littlejohn and Deitrich, 1972; Dutton *et al*., 1981; Stone and Wilson, 1982), further validating our measurements. The minor discrepancy may be due to differences in experimental protocols, tissue handling, or the use of incubation media and chemical treatments in some of those studies. Furthermore, compared with anatomically adjacent tissues, the iris is more hydrated. Namely, the cornea has been reported to contain approximately 80% water content (Monti *et al*., 2002), while the crystalline lens of mature rabbit has 50% and 69% water content in the nucleus and cortex, respectively (Hockwin *et al*., 1978), and the sclera 71% (Boubriak *et al*., 2000). The iris is the most dynamic tissue in the eye, and its high water content suggests it can actively absorb and release fluid into the anterior chamber, which could be a significant factor in understanding anterior chamber homeostasis and glaucoma pathology.

As an illustrative quantitative example, iris contractions can significantly change human iris volume (Quigley *et al*., 2009; Mak, Xu and Leung, 2013; Seager, Jefferys and Quigley, 2014; Liao *et al*., 2024). Some reports indicate that during light-to-dark dilation, the iris loses 1.8%–2.8% volume per mm change in pupil diameter (Mak, Xu and Leung, 2013; Liao *et al*., 2024), while pharmacologically induced dilation can reduce iris volume as high as 4.4% per mm change in pupil diameter (Mak, Xu and Leung, 2013). Considering that a healthy iris has a volume of c. 40 µL in photopic light conditions (Mak, Xu and Leung, 2013), iris dilation can release 0.6–1.6 µL per mm change in pupil diameter, or 3.6–9.6 µL—assuming the full range of pupil dilation from 2 mm to 8 mm and 88% water content based on our results. As a reference, the human eye produces 2.2–3.1 µL/min of aqueous humor (McLaren, 2009), which indicates that the fast (almost instantaneous) iris contractions can impact aqueous humor dynamics. While volume loss during dilation is not universal (Seager, Jefferys and Quigley, 2014; Liao *et al*., 2024), disease state, contraction mechanism, and other anatomical factors can influence the magnitude of these estimates, and hydraulic permeability and the speed of iris contraction can influence the rate at which these changes occur. Therefore, further investigation is needed on the connection between iris contractions and aqueous humor dynamics.

One novel aspect of our data is the focus on sex differences, which are particularly important for PACG studies. In humans, women are at higher risk of developing PACG (Vijaya et al., 2006; Cheng et al., 2013; Zhang et al., 2021), and some experimental evidence also suggests distinct biomechanical properties between male and female irises, with female murine irides being stiffer than males (Lee *et al*., 2021). Our study found that the male iris has slightly higher water content, which in theory can affect tissue stiffness (Screen *et al*., 2006; Safa *et al*., 2017). Although we noted a statistically significant result, the practical impact of this small difference (c. 1.5%) is likely minimal. As a related technical note, we also conducted a gross morphology analysis to assess the potential confounding effects of differential postmortem dilation in our measurements, which may arise from differences in muscular tone or biomechanical properties. However, our results showed no differences in limbal or pupil diameter, or in their ratio, between sexes (Figure 2). These findings add additional insight into using rabbits as an animal model for studying iris biomechanics. Whereas, although natural development of PACG in rabbits is not common (Jeong *et al*., 2005), rabbits, due to the shape of their pupil and the size of the iris, are a highly suitable model for iris biomechanics (Yamaji *et al*., 2003; Zhang *et al*., 2014; Li *et al*., 2021, 2023); therefore, the potential equivalence of the male and female biomechanical properties in rabbits would help reduce the need for accounting for sex effects in the procurement, reducing overall cost and study complexity. However, one cannot rule out other potential sex-dependent biomechanical differences, which merit further investigation.

Furthermore, inspired by the ubiquitous use of PBS in experimental biomechanical tests, we specifically tested the effect of PBS buffer exposure on iris water content, (Safa *et al*., 2017; Bloom, Lee and Elliott, 2021). Buffer solutions are typically used in these tests to maintain the water content of the tissue *ex vivo*; however, they may affect the biomechanical properties of the tissue through alteration of the water content or ionic interaction of the buffer solutes with the fixed negative charge that naturally exists in many soft tissues due to the glycosaminoglycans (GAGs) content of the tissues (Lujan *et al*., 2009). Our results showed minimal effect of PBS on water content (Figure 5), with female eyes showing a small change compared to their controls, although typically an increase in water content is expected after exposure to PBS. This effect can be explained by the similar osmolality of PBS and aqueous humor, which are 280 ± 2.6 mOsm/kg (Hocking *et al*., 2012) and 313 ± 14 mOsm/kg^1^ (Huang *et al*., 2021), respectively. An alternative explanation could be regional changes in iris permeability, such as in its epithelial layers, which may cause a heterogeneous distribution of water content in the tissue that our measurement technique could not detect.

One limitation of our study was the exclusive use of cadaveric eyes at a fixed level of postmortem dilation; thus, similar experiments may be needed on irides at varying levels of contraction, which can impact water content as discussed previously. Nevertheless, our analysis indicated a similar state of postmortem dilation (Figure 2), which would reduce the likelihood that differential dilation affected male and female samples. However, further experimentation using pharmacological induction of iris contraction with various miotic and mydriatic agents (Aptel and Denis, 2010; Li *et al*., 2018) could provide insight into the effects of iris contraction and its water content. Such experiments may need to be performed shortly after animal termination—often within a few hours (Whitcomb *et al*., 2009)—to preserve iris smooth muscle contractility and may pose practical challenges.

Furthermore, we conducted gross morphology measurements using two-dimensional imaging (Figure 1), overlooking potential three-dimensional effects; three-dimensional AS-OCT imaging may be required to further validate the accuracy of our gross morphology results. Moreover, our investigation included only mature rabbits, so our findings may not be generalized to other ages or species. As a result, a similar detailed analysis is warranted using, for example, geriatric human eyes, which are the primary population group of PACG (Zhang *et al*., 2021). Additionally, our hydration findings include the water content of rabbit irides with the ciliary body attached, which may confound results because these two uveal tract tissues may differ compositionally. Because removing rabbit ciliary muscles was not possible without inducing damage and unwanted mechanical strain during sample preparation, we retained the ciliary body in our samples, which is a limitation. Finally, although we took every possible measure to minimize the effects of sample preparation, dissecting the iris and manually blotting to remove loose aqueous humor were inevitable and could have caused variations in water content that were not measurable in the current study; however, we used the same protocol across tests, and this effect is unlikely to have significantly affected the interpretation of our findings.

In conclusion, this study provided a detailed experimental measurement of iris water content and sex differences in rabbit irides. We noted that female irides, despite having a similar gross morphology, had slightly lower water content. Additionally, 20 minutes of PBS exposure had minimal to no effect on iris water content. Our results improve understanding of iris physiology and can be used to evaluate the tissue’s biomechanical response under mechanical loading, enabling the creation of physiologically relevant computational models to better assess the role of iris biomechanics in PACG.

## Conflicts of interest

The authors declare no conflicts of interest.

## Acknowledgments

We acknowledge NIH/NEI (R00EY035360 [BNS]) as the funding source for this study. The content is solely the responsibility of the authors and does not necessarily represent the official views of the NIH.

## Footnotes

1 Individual data points are extracted by digitization of Fig. 2D of Huang *et al*., 2021 using WebPlotDigitizer (https://automeris.io/).

## Notes

### Competing Interest Statement

The authors have declared no competing interest.

