## Supplementary Document for "Experimental Assessment of Iris Water Content and Sex Differences in Mature Rabbits"

### Figure S1

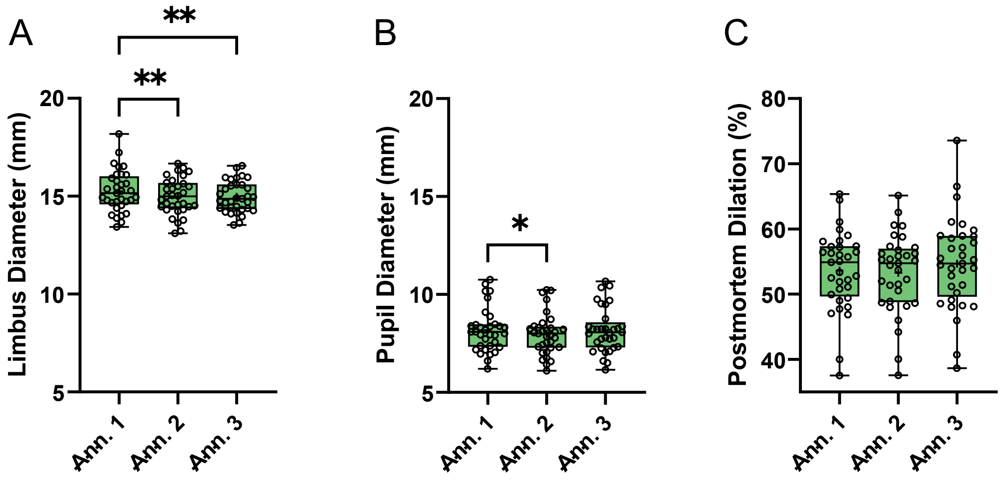


**Figure S1: Gross morphology analysis for each annotator (abbreviated as Ann.) shown for (A) equivalent limbus diameter, (B) equivalent pupil diameter, and (C) postmortem dilation ratio. One-way repeated-measures ANOVA followed by Tukey's multiple comparisons test despite not showing a difference for the equivalent pupil diameter (B; p = 0.095) and postmortem ratio (C; p = 0.100), it indicated some differences between annotators for the equivalent limbus diameters (p < 0.001); therefore we averaged the measurements for the final analysis. The multiple comparisons indicated difference in both equivalent limbus diameter (A) and pupil diameter (B). Although some of the inter-annotator differences were statistically significant, the overall difference was modest. This variability may be partially explained by blurry transitions at the margins between the iris, pupil, and sclera (Figure 1). Data are shown as individual points with box and whiskers as before. Multiple comparison results shown with *p < 0.05 and **p < 0.01**
